# Angiopoietin like protein 3 regulates low-density lipoprotein transport through aortic endothelial cells via endothelial lipase

**DOI:** 10.1101/2025.09.30.679508

**Authors:** Yen Way Isabell Trinh, Grigorios Panteloglou, Evelina Voloviceva, Museer A. Lone, Stephanie Bernhard, Eveline Schlumpf, Kathrin Frey, Lucia Rohrer, Mary Juliette Melo Gonzales, Thomas Lutz, Saša Frank, Arnold von Eckardstein, Jerome Robert

**Affiliations:** Institute of Clinical Chemistry, University Hospital of Zurich, University of Zurich, Zurich, Switzerland; Institute of Veterinary Physiology, Vetsuisse Faculty University of Zurich, Zurich Switzerland; Gottfried Schatz Research Centre, Molecular Biology and Biochemistry, Medical University of Graz, 8010 Graz, Austria

## Abstract

Angiopoietin-like protein 3 (ANGPTL3) inhibits endothelial lipase (EL, encoded by LIPG) and lipoprotein lipase (LPL) and has emerged as a promising target for lipid-lowering therapies. While ANGPTL3 inhibition lowers circulating triglycerides and low-density lipoprotein cholesterol (LDL-C), its impact on atherosclerotic cardiovascular disease (ASCVD) remains uncertain. This study investigates the roles of ANGPTL3 and EL in the trans-endothelial transport of LDL and high-density lipoprotein (HDL), an early step in atherogenesis. Using primary human aortic endothelial cells (HAEC), we demonstrate that EL promotes binding, uptake, and transcytosis of LDL and HDL, with its catalytic activity being essential for transport but not for surface association. ANGPTL3 selectively reduced the transport of LDL but not HDL through HAEC. Mechanistically, EL and scavenger receptor BI (SR-BI) act sequentially to mediate LDL uptake, but independently of each other towards HDL uptake. *Ex vivo,* ANGPTL3 reduced LDL accumulation within bovine aorta. Interestingly, ANGPTL4 had no measurable effect on lipoprotein transport, while ANGPTL8 modestly inhibited both LDL and HDL association with endothelial cells. Our findings provide evidence that ANGPTL3 inhibits EL-mediated transendothelial LDL transport. Opposing effects of ANGPTL3 inhibition on LDL-C concentration and LDL entry into the arterial wall may explain why recent population studies did not reveal any genetically causal association of ANGPTL3 and LIPG with ASCVD risk.

## 1. Introduction

The findings of hypolipidemia in human carriers of loss of function (LOF) mutations in the angiopoietin like protein 3 (*ANGPTL3*) gene have positioned ANGPTL3 as an attractive target for the treatment of severe hypertriglyceridemia and hypercholesterolemia ^1–5^. Indeed, monoclonal human ANGPTL3-blocking antibodies (Evanicumab) lowered plasma levels of triglycerides, total-cholesterol, low-density lipoprotein-cholesterol (LDL-C), and high-density lipoprotein-cholesterol (HDL-C) in patients with familial hypercholesterolemia during phase-2 and phase-3 clinical trials— independently of LDL receptor (LDLR) activity ^6–8^. Recently, another monoclonal antibody targeting ANGPTL3 successfully reduced LDL-C levels in patients with suboptimally controlled hyperlipidemia in a phase-2 clinical trial ^9^. An antisense oligonucleotide (ASO, Vupanorsen), which inhibits ANGPTL3 protein synthesis, demonstrated similar lipid-lowering effects in phase-2 clinical trials, though development was halted due to increased hepatic fat fraction and elevated level of alanine aminotransferase (ALT) and aspartate aminotransferase (AST) ^10–12^. Most recent phase-2 trials investigating RNA inhibition strategies (Zodasiran and Solbinsiran) have also shown promise in reducing blood lipid levels in patients with mixed hyperlipidemia ^13–15^.

ANGPTL3 is a glycoprotein secreted by the liver that acts as an inhibitor of lipoprotein lipase (LPL) and endothelial lipase (EL). However studies in mice lacking both EL and LDLR, (*Lipg^-/-^*ldlr^-/-^*) revealed that the LDL-C-lowering effect of ANGPTL3 inhibitors depends on the presence of EL. The disinhibition of EL appears to be the mechanism by which ANGPTL3-targeting antibodies, ASOs, and siRNAs lower plasma HDL-C levels as well. LOF variants of LIPG increase plasma levels of HDL-C and also improve HDL functionality ^16^. A combined meta-analysis of the Rotterdam, GiraFH and premature atherosclerosis (PAS) cohorts investigating eight different *LIPG* LOF mutations, showed a reduced risk of atherosclerotic cardiovascular disease (ASCVD) compared to non-carriers ^16^. However Mendelian randomization analyses specifically investigating the *LIPG396*Ser allele, which is associated with higher HDL-C, failed to find any significant association with a history of myocardial infarction ^17^. These findings challenge the notion that inhibiting ANGPTL3 and -thereby enhancing EL function-would universally reduce ASCVD risk. In fact, recent Mendelian randomization studies investigating LOF variants mutations that reduce ANGPTL3 levels failed to show any impact on ASCVD risk, despite lowering LDL-C and triglycerides ^18,19^. Therefore, the hypolipidemic and hence anti-atherogenic effects of lower ANGPTL3 activity and higher EL activity in the liver may be counteracted by other pro-atherogenic effects.

The transendothelial transport of lipoproteins into the vascular wall has caught increasing attention as an important pathogenic mechanism in ASCVD ^20^. Caveolin-1, scavenger receptor BI (SR-BI), and activin-like kinase 1 (ALK1) facilitate the transport of LDL ^20^. Our lab previously identified EL as a limiting factor in transendothelial HDL transport in addition to SR-BI, ATP binding cassette transporters ABCA1 and ABCG1, and the ecto-F1F0-ATPase/purinergic receptor axis, ^21,22^. We therefore investigated here, if EL and ANGPTL3 also influence the transport of LDL through the endothelium.

## 2. Material and methods

### 2.1 Cells

Primary human aortic endothelial cells (HAEC; passages 2–7, PromoCell, Germany, and Lonza, Switzerland) were maintained in endothelial growth medium (EGM-2; Lonza, Switzerland) supplemented with SingleQuots™ (Lonza) and 10% fetal bovine serum (FBS; Sigma-Aldrich, Switzerland). The endothelial cell line EA.hy926 (ATCC, USA) were cultured in Dulbecco’s Modified Eagle Medium (DMEM; Sigma-Aldrich) containing L-glutamine and 10% FBS.

### 2.2 Lipoproteins

LDL and HDL were isolated from normolipidemic individuals using KBr gradient ultracentrifugation as described previously ^23,24^. The purity of the isolated lipoproteins was assessed by SDS-PAGE followed by Coomassie Blue staining. LDL and HDL were radiolabeled with I-125 (Hartmann, Germany) using a modified version of McFarlane’s protocol adapted for lipoproteins^23,24^. Specific activity was determined using a Wizard2 gamma counter (PerkinElmer, USA) and ranged between 150–500 cpm per ng of total lipoprotein protein.

Alternatively, LDL and HDL were fluorescently labeled with Atto dyes (AttoTec, Germany) following the manufacturer’s instructions. Briefly, 1 mg of total lipoprotein protein was diluted to a concentration of 2 mg/mL in phosphate-buffered saline (PBS; 137 mM NaCl, 2.7 mM KCl, 10 mM Na_2_HPO_4_, 1.8 mM KH_2_PO_4_) and incubated with 10 µg of NHS-ester Atto dye (488 or 655). Labeling efficiency was enhanced by adding sodium bicarbonate (1 M, pH 9) to a final concentration of 0.1 M. After 1 hour in the dark at room temperature with constant gentle rotation, fluorescently labeled lipoproteins were separated from free dye using PD-10 desalting columns (Cytiva, Switzerland) pre-equilibrated with PBS.

Both radiolabeled and fluorescently labeled lipoproteins were stored at 4 °C for a maximum of one month.

### 2.3 LIPG knockdown by siRNA transfections

HAEC and Huh-7 cells were reverse transfected with siRNA targeting *LIPG* (catalog no. M-009601-01-0005, Horizon Discovery Dharmacon™, USA), *ANGPTL4* (catalog no. M-007807-02-0010, Horizon Discovery Dharmacon™), or a non-targeting control sequence (catalog no. D-001810-10-50, Horizon Discovery Dharmacon™). Briefly, trypsinized cells in complete growth medium were added at an 8:2 ratio to a mixture of siRNA (final concentration 10 nM) and Lipofectamine™ RNAiMAX (final dilution 1:1000; Invitrogen, Thermo Fisher Scientific, USA) in Opti-MEM™ (Gibco, Thermo Fisher Scientific). Cells were then cultured for 72 hours without a media change prior to experimentation.

### 2.4 EL overexpression using adenoviruses

HAEC and EA.hy926 cells were infected with adenovirus (MOI 50) encoding either wild-type human EL (EL_WT), a catalytically inactive EL mutant containing the D192N substitution (EL_MUT), or an empty vector control (Ad_vec). Cells were cultured with the adenovirus for 48 hours prior to experimentation. Adenoviral vectors were prepared as previously described ^25^.

### 2.5 Stable SCARB1 knockdown using lentivirus-based shRNA

Stable EA.hy926 cells lacking SR-BI were generated as previously described ^26^. Briefly, EA.hy926 cells were transduced with lentivirus shuttle plasmid for shRNA targeting human *SCARB1* (TRCN0000056966, pLKO.1, ETH NEXUS Personalized Health technology, Switzerland) or a control lentivirus shuttle plasmid for non-targeting scramble shRNA (Addgene plasmid #18649, a gift from David Sabatini; (Addgene plasmid #1864; http://n2t.net/addgene:1864; RRID:Addgene_1864) both produced in HEK 293T cells. Transductions were carried out in the presence of polybrene (8 μg/mL; Sigma-Aldrich) to enhance infection efficiency. After transduction, cells were selected with puromycin (1 μg/mL; Sigma-Aldrich) for four passages prior experimentation. The efficiency of *SCARB1* silencing was assessed by RT-qPCR and Western blotting.

### 2.6 Pharmacological inhibition of EL

HAEC were treated with 1 μU/mL of heparinase III (Sigma-Aldrich) in DMEM supplemented with 0.2% bovine serum albumin (BSA; Sigma-Aldrich) for 30 minutes prior to the assays. In other experiments, 10 μM tetrahydrolipstatin (THL; Sigma-Aldrich, Switzerland), 1 μg/mL recombinant human ANGPTL3 (Abcam, USA), or 1 μg/mL recombinant human ANGPTL8 (R&D Systems, USA) were added to HAECs 30 minutes before the assay but unlike heparinase III, maintained in the medium throughout the assay.

### 2.7 Lipoprotein binding

One hundred thousand cells per well were seeded in a 24-well plate in complete growth medium. On the day of the assay, the medium was replaced with ice-cold DMEM supplemented with HEPES (Sigma-Aldrich) and 0.2% BSA. After a 30-minute incubation on ice, the medium was removed, and cells were incubated with 10 μg/mL of I^125^–labeled lipoproteins in DMEM containing HEPES and 0.2% BSA, in the absence or presence of a 40-fold excess of the respective unlabeled lipoproteins. Cells were maintained at 4 °C for 1 hour before removing the assay medium and washing twice with 50 mM Tris-HCl (pH 7.4), 150 mM NaCl, 0.02% NaN_3_, and 0.2% BSA, followed by a final wash with PBS containing 0.1 mM CaCl_2_ and 1 mM MgCl_2_. Cells were then lysed in 0.2 M NaOH for at least 30 minutes before measuring radioactivity using a Wizard2 gamma counter (PerkinElmer, USA). Counts per minute (cpm) were normalized to cellular protein content, determined using the DC protein assay (Bio-Rad Laboratories, USA). Specific binding of both LDL and HDL was calculated by subtracting the cpm measured in the presence of a 40-fold excess of unlabeled lipoproteins from the cpm measured with labeled lipoproteins alone.

### 2.8 Lipoprotein association

One hundred thousand cells per well were seeded in 24-well plates and cultured to confluence for 2 to 3 days. On the day of the assay, cells were incubated with 10 μg/mL of I^125^–labeled lipoprotein in DMEM supplemented with HEPES and 0.2% BSA, in the absence or presence of a 40-fold excess of unlabeled LDL or HDL. After an hour incubation at 37 °C, cells were processed as described for the binding assay.

### 2.9 Lipoprotein transport

One hundred thousand HAEC were seeded onto Transwell inserts (pore size 0.3 μm; Corning, USA) placed in 24-well plates and cultured to confluence for 3 days. For experiments involving viral infection, cells were infected 24 hours after seeding and maintained in culture for an additional 2 days. On the day of the assay, 10 μg/mL of I^125^–labeled lipoprotein in DMEM supplemented with HEPES and 0.2% BSA, with or without a 40-fold excess of the respective unlabeled lipoprotein, was added to the upper chamber of the Transwell. After a 1-hour incubation at 37 °C, media from the lower chamber was collected and counted. Cells were then processed as described for the binding assay.

### 2.10 Lipoprotein uptake

Three hundred thousand cells were incubated at the indicated temperature with 40 μg/mL Atto488-LDL or 40 μg/mL Atto655-HDL in DMEM supplemented with 0.2% BSA, either in the presence (non-specific uptake) or absence (total uptake) of a 40-fold excess of the respective unlabeled lipoproteins. After 1 hour at the indicated temperature, cells were extensively washed with PBS and detached using Accutase® (Sigma-Aldrich) for 5 minutes at 37 °C. Following detachment and washing with PBS, samples were incubated with 2 μg/mL propidium iodide (PI; Fluka, Switzerland) to exclude dead cells. Sample acquisition was performed on a BD LSR II Fortessa flow cytometer (BD Biosciences, USA) using BD FACSDiva™ software. Cell debris was excluded by gating on forward scatter area versus side scatter area (FSC-A/SSC-A), and doublets were excluded via FSC-A/FSC-H gating. Live cells were identified by selecting PI-negative events through PI-A/FSC-H gating. To correct for spectral overlap between fluorophores, unstained and single-stained controls were acquired and compensation was applied using FACSDiva™ software.

For each condition, approximately 2 × 10^4^ events within the final live cell gate were recorded and used for analysis. Data were analyzed using FlowJo version 10 (FlowJo LLC, USA). The median fluorescence intensity (MFI) of each population was used for comparison across conditions, and specific uptake was calculated as the difference between total and non-specific uptake.

### 2.11 Lipoprotein degradation

HAEC were seeded at 1 × 10^5^ cells per well in 24-well plates and cultured to confluence for 2 to 3 days. On the day of the assay, cells were incubated with 10 μg/mL of I^125^–labeled LDL or HDL in DMEM supplemented with HEPES and 0.2% BSA, either in the absence (total uptake) or presence (non-specific uptake) of a 40-fold excess of the respective unlabeled lipoprotein. After 4 hours of incubation, the assay medium was collected and ice-cold trichloroacetic acid (TCA, Sigma-Aldrich) was added to a final concentration of 12% and mixed thoroughly. Following a 30-minute incubation at 4 °C, samples were centrifuged at 2000 × g for 10 minutes at 4 °C. The supernatant was transferred to new tubes containing sodium iodide (NaI; final concentration 0.4%, Sigma-Aldrich) and vortexed for 5 seconds. After 5 minutes at room temperature, hydrogen peroxide (H_2_O_2_ Sigma-Aldrich) was added to a final concentration of 1.1%. Five minutes later, chloroform was added and the upper aqueous phase containing the lipoprotein degradation products was collected and measured using a Wizard2 gamma counter to quantify degradation.

The cells in the assay plate were washed twice with Tris-BSA and once with PBS containing calcium and magnesium, lysed with NaOH, and the associated radioactivity was also measured using the Wizard2 gamma counter. The percentage of degradation relative to association was calculated as:

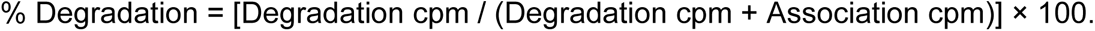

### 2.12 Determination of Lipoprotein size

Lipoprotein size before and after transendothelial transport was determined using size exclusion Fast Protein Liquid Chromatography (FPLC; Cytiva, USA). Briefly, HAEC overexpressing wild-type EL (WT_EL) or transduced with an empty vector were cultured on Transwell inserts as described above. Forty-eight hours post adenoviral infection, 50 μg/mL of I^125^–labeled HDL or LDL were added to the upper chamber and incubated for 1 hour at 37 °C. Media from the lower chamber was collected and mixed with saturated sucrose (2:1 ratio) and 1 mg of unlabeled lipoprotein to protect the radiolabeled lipoproteins from oxidation and degradation. For size separation, 500 μL of the collected material was loaded onto a HiLoad 16/600 Superdex 200 pg size exclusion column (Cytiva). Isocratic elution was performed at 4 °C using 150 mM NaCl and 10 mM Tris (pH 7.5) at a constant flow rate of 1 mL/min. Fractionation began immediately after sample injection, with 1 mL fractions collected throughout the run. Each fraction was measured using a gamma counter to assess the distribution of radioiodinated particles transported across the endothelial layer. Empty Transwell inserts (lacking cells) served as control conditions to monitor for spontaneous particle modification.

### 2.13 Untargeted lipidomics by LC-MS/MS analysis

Two and a half million EA.hy926 cells were infected with adenovirus encoding EL_WT or EL_MUT. After 48 hours, the medium was collected and centrifuged to remove dead cells and debris. LDL or HDL (100 µg/ml) from three individual donors were then incubated in 1 ml of the medium. After 4 hours, the collected media fractions containing lipoproteins were frozen at -80°C and lyophilized. Dried fractions were then extracted with 1 ml Methanol/MTBE/Chloroform (MMC) (4/3/3) containing 50 mg/L of Butylated hydroxytoluene (BHT) as described here ^27^. Extraction solvents were supplemented with SPLASH lipidoMIX internal standard (Avanti Polar Lipids, Alabaster, AL) and in-house sphingolipidomics ^27^ standard mix. Briefly, lipids were extracted continuously on a Thermomixer (Eppendorf) at 37°C (1400 rpm, 60 minutes). Extracts were then centrifuged at 16000 g /10 min /RT). The single-phase supernatant was collected in fresh tubes and dried under N_2._ Dried lipids were finally dissolved in 100 μl methanol on a Thermomixer (650 rpm, 24°C, 60 minutes), spun (16000 g /10 min/RT) and transferred to MS vials. Untargeted lipid analysis was performed from 10 µl of the injected extracts on a high-resolution Q-Exactive MS analyzer (Thermo Scientific) after lipids were separated by liquid chromatography. Lipids were separated using a C18 Acquity UPLC column (150 mm x 2.1 mm, 1.7 µm particle size) and a Transcend UHPLC pump (Thermo Fisher Scientific). Liquid chromatography was performed with solvents, acetonitrile:water (6:4) with 10 mM ammonium acetate and 0.1 % formic acid, B) isopropanol:acetonitrile (9:1) with 10 mM ammonium acetate and 0.1 % formic acid at a flow rate of 0.260 ml/min exactly as described ^27^. MS2 fragmentation was based on data dependent acquisition (DDA). Tracefinder 5.1 (Thermo Fischer Scientific) was used for lipid identification at an accuracy of 5 ppm from the predicted mass at a resolving power of 70’000 at 200 m/z.

### 2.12 Quantification of mRNA

Total RNA was isolated from 4 × 10^5^ cells HAEC using either the RNA isolation kit (Macherey Nagel, Germany) or the NucleoSpin™ Mini Kit for RNA Purification (Quiagen, Germany) according to manufacturer’s instructions. Genomic DNA was removed using on-column DNAse treatment. Equal amount of RNA (0.5-2 μg) were reverse transcribed using random hexamers and the RevertAid First Strand cDNA Synthesis enzyme (Thermo Fischer Scientific). Real-time quantitative polymerase chain reaction (RT-qPCR) was performed using LightCycler^®^ 480 SYBR Green I Master reagent (Roche) on a Light Cycler 480-II system (Roche) to quantify transcript levels. Specific primer: *LIPG* (fwd: TTCACGGATGGACGATGAGC, rev: GGAGCCAGTCAACCACAACT), *SCARB1* (fwd: CTGTGGGTGAGATCATGTGG, rev: GCCAGAAGTCAACCTTGCTC), LDLR (fwd: AAGGACACAGCACACAACCA, rev: CATTTCCTCTGCCAGCAACG), *ALK1* (fwd: CCTGTGGCATGTCCGACG, rev: TAGCGGCCTTTTCCCCCCACACA), *ABCG1* (fwd: GTCGCTCCATCATTTGCACC, rev: ATTGCAGACTTTTCCCCGGT) and *GAPDH* (fwd: CCCATGTTCGTCATGGGTGT, rev: TGGTCATGAGTCCTTCCACGATA ). Relative transcript expressions was calculated using the 2^−ΔΔCt^ method, with *GAPDH* serving as the internal reference.

### 2.13 Sodium dodecyl sulfate (SDS) polyacrylamide gel electrophoresis (PAGE)

HAEC were lysed in ice-cold radioimmunoprecipitation assay buffer (RIPA, 5 mM EDTA, 50 mM NaCl, 10 mM sodium pyrophosphate, 50 mM NaF, 1% NP-40 alternative, pH 7.4) supplemented with cOmplete™ Protease Inhibitor Cocktail (Roche). Lysates were centrifuged at 14,000 rpm for 20 minutes at 4 °C, and total protein concentrations in the supernatants were quantified. Equal amounts of protein (20–30 µg) were separated by SDS-PAGE and transferred to polyvinylidene fluoride (PVDF) membranes (AmershamTM hybind® P, Sigma-Aldrich). Following transfer, membranes were stained with Ponceau S to verify protein transfer, then blocked in 5% skim milk in PBS containing 0.1% Tween-20 (PBST) for at least 30 minutes. Membranes were then incubated with primary antibodies against EL (NB400-118, 1:1000, Novus), SR-BI (NB400-101, 1:1000, Novus), ALK1 (ab21870, 1:1000, ABCAM), LDLR (ab52818, 1:1000, ABCAM), and GAPDH (Ab9484, 1:1000, ABCAM) for 2 hours at room temperature or overnight at 4 °C. After three washes in PBST, membranes were incubated with HRP-conjugated anti-mouse or anti-rabbit secondary antibodies (1:10,000, P0260 and P0448, respectively; Dako, Agilent Pathology Solutions) in PBST containing 5% milk for 1 hour at room temperature. Membranes were washed three times with PBST for 5 minutes each and developed using the WesternBright Sirius Chemiluminescent Detection Kit (K-12043-D20, Advansta). Signal detection was performed using the Fusion FX imaging system (Vilber Lourmat, France).

### 2.13 Phospholipase assay

EL activity was measured using the Phospholipase A2 PED-A1 kit (Invitrogen, Thermo Fischer Scientific) according to the manufacturer’s instructions. Briefly, 2.5 × 10^6^ cells were seeded in 10 cm culture dishes and grown to confluence (48–72 hours). Culture media were collected and centrifuged at 300 × g for 5 minutes at room temperature. Then, 50 µL of the clarified media were incubated with 50 µL of the lipase substrate mix in the presence or absence of 1 µg/mL ANGPTL3 at 37 °C. Fluorescence was measured at the indicated time points using a plate reader (excitation: 450 nm; emission: 515 nm; Tecan Trading AG, Switzerland). The time course of lipase activity was plotted, and the area under the curve (AUC) was calculated using Prism 10 (GraphPad Software, USA)

### 2.14 Immuno-staining of EL in cells and bovine aorta

One hundred thousand cells were seeded on glass coverslips 24 hours prior to infection with adenoviruses encoding wild-type EL (EL_WT), mutant EL (EL_MUT), or empty vector (Ad_vec). After 48 hours, cells were extensively washed with PBS and fixed with 4% paraformaldehyde (PFA) at room temperature for 20-45 minutes. Cells were then washed once with 0.5 M Tris-HCl (pH 8.0) and twice with PBS. Following 5 minutes of permeabilization with 0.5% Triton X-100 in PBS and additional PBS washes, cells were blocked with 5% donkey serum and 1% BSA in PBS for at least 1 hour. Cells were then incubated overnight at 4 °C with primary anti-EL antibody (Ab100987, 1:50 ABCAM). After extensive PBS washes, cells were incubated for 1 hour at room temperature with anti-rabbit secondary antibodies (Invitrogen, Therma Fischer Scientific, 1:600) and 1 ng/mL DAPI (Sigma Aldrich). After final PBS washes, coverslips were mounted in Vectashield (Vector Laboratories, USA) and imaged using a SP8 confocal microscope (Leica, Germany).

Bovine aortae were obtained from the slaughterhouse of the Vetsuisse Faculty University of Zurich. Immediately after slaughter, a segment of the aortic arch (5-10 cm) was placed in DMEM containing 10% FBS and antibiotics/antimycotics (100 U/mL penicillin, 100 µg/mL streptomycin, 0.25 µg/mL amphotericin B; Gibco, Thermo Fisher Scientific). Tissues were kept on ice for a maximum of 4 hours. Aortic segments were then washed three times with PBS and fixed in 4% PFA in PBS at 4 °C for 48 hours. Fixed tissues were washed once for 10 minutes with 0.5 M Tris-HCl (pH 8.0) and twice with PBS. Tissues were then incubated in 20% sucrose in PBS for 48 hours at 4 °C. Sections were cut at 20 µm using a cryostat (Leica, Germany) and stored at −80 °C until further processing. For immunostaining, aortic sections were brought to room temperature for 10 minutes and rehydrated twice in PBS (10 minutes each). Sections were permeabilized with 0.5% Triton X-100 in PBS for 5 minutes, followed by three PBS washes. After blocking with 5% donkey serum (Sigma-Aldrich) and 1% BSA in PBS for at least 60 minutes, sections were incubated overnight at 4 °C with primary antibodies against EL (nb400-118, 1:50, Novus). After three 5-minute PBS washes at room temperature, sections were incubated for 1 hour with anti-rabbit and anti-mouse secondary antibodies (1:600; Invitrogen, Thermo Fischer Scientific) diluted in blocking solution, along with DAPI (1 ng/mL). Sections were washed three additional times with PBS (5 minutes each), mounted in Vectashield, and imaged using a SP8 inverted confocal microscope. Microscopy images were used for illustrative purposes only and were not quantified.

### 2.16 Ex vivo LDL and HDL uptake

Fresh bovine aortae (n = 6) were collected as described above. From the aortic arch, 5 mm diameter tissue punches were obtained using a biopsy device (Kai Medical, Japan) and incubated overnight in DMEM supplemented with 10% FBS and antibiotics/antimycotics in a humidified incubator at 37 °C with 5% CO₂. The following day, aortic pieces were embedded in 2% agarose in PBS within a 12-well plate, with the endothelial surface facing upward and exposed above the gel. Tissues were then incubated with 10 µg/mL of I^125^–labeled lipoproteins for 2 hours. After incubation, aortic pieces were washed three times with Tris-HCl containing 0.2% BSA, mechanically removed from the agarose, and further washed three times with PBS containing calcium and magnesium. Samples were then transferred into tubes containing 0.2 mM NaOH, and radioactivity was measured using a Wizard² gamma counter as previously described.

Alternatively, bovine aortic tissues were incubated with 100 µg/mL of fluorescently labeled (Atto) lipoproteins for 2 hours. At the end of the incubation period, tissues were placed on ice and washed five times with ice-cold PBS containing calcium and magnesium. Pieces were then fixed in 4% PFA for 24 hours at 4 °C, washed once for 10 minutes with 0.5 M Tris-HCl (pH 8.0), and twice with PBS. After fixation, tissues were incubated in 20% sucrose in PBS for an additional 24 hours at 4 °C. Aortic pieces were then embedded in Tissue-Tek O.C.T. (Qiagen), sectioned, stained, and imaged as described above.

### 2.17 Statistics

For all analyses, except RT-qPCR, raw linear data were first log-transformed and analyzed using a blocked Student’s *t*-test or one-way ANOVA with Dunnett’s post hoc test, with "Experiment" used as the blocking factor. For RT-qPCR analyses, 2^−ΔΔCt^ values were analyzed using the same statistical tests. Data were obtained from at least three independent experiments and are graphically represented as biological replicates with mean ± standard deviation (SD). Experiments involving bovine aortic tissues were analyzed using a paired Student’s *t*-test after confirming normality with the Shapiro–Wilk test. Statistical analyses were performed using SPSS Statistics 25 (IBM, Chicago, IL, USA) or Prism 8 (GraphPad Software, USA), with *p* ≤ 0.05 considered statistically significant. All graphs were generated in Prism 8, with control (vehicle, non-coding or empty vector) conditions normalized to 100%. Statistical analysis of the lipidomics data was performed using the MetaboAnalyst 6.0 (https://doi.org/10.1093/nar/gkae253) platform (One Factor Statistical Analysis module). The lipid concentration table was uploaded as the input file. Left-censored data estimation was used for missing value imputation and data were log10-transformed. For the volcano plots, paired analysis was performed (Fold Change threshold = 2.0, raw p-value threshold = 0.05). Result table was downloaded and used to visualize the volcano plots in GraphPad Prism 8.

## 3 Results

### 3.1 Inhibition of endothelial lipase reduces both LDL and HDL binding and uptake by endothelial cells

We previously reported that EL regulates HDL transport through bovine aortic endothelial cells ^22^. Since EL binds and hydrolyzes phospholipids in both HDL and LDL in the plasma ^28^, we investigated whether EL also limits LDL transport through aortic endothelial cells. First, we used pharmacological inhibitors of EL and measured both LDL and HDL association with HAEC. Lipoprotein association, performed at 37°C, records both the binding to and uptake by endothelial cells. A 20-minute treatment with heparinase, which cuts the heparin-sulfate proteoglycans (HSPG) where EL binds on the endothelial cell surface ^28^, significantly reduced both LDL and HDL association with HAEC (***Figure 1A-B***). Moreover, inhibition of EL with the lipase inhibitor THL also significantly reduced LDL and HDL association (***Figure 1C-D***).

**Figure 1.**
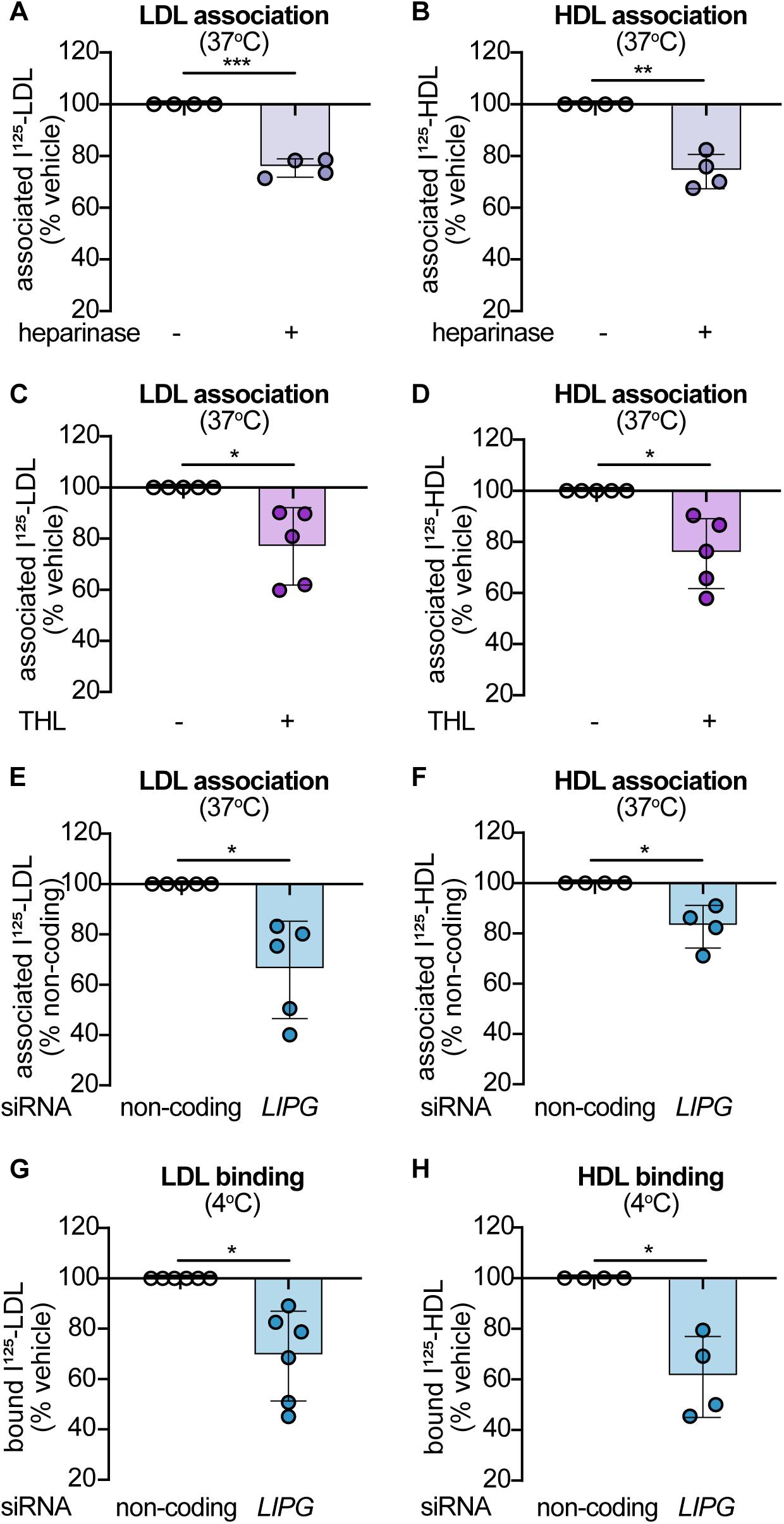
Inhibition of endothelial lipase reduces LDL and HDL association with human aortic endothelial cells. **A** through **D)** Confluent HAEC were incubated with 1 μU/mL heparinase (A-B) or 10 μM THL (C-D). After 30 minutes, the cells were incubated with 10 μg/mL of I¹²⁵-LDL (A and C) or I¹²⁵-HDL (B and D) for 60 minutes at 37 °C, followed by extensive washing and cell lysis. Radioactivity was measured using a γ-counter. **E** and **F)** Three days after siRNA transfection against *LIPG*, I¹²⁵-LDL (**E**) or I¹²⁵-HDL (**F**) associations were measured in confluent HAEC as described above. **G** and **H**) Three days after transfection against LIPG, confluent HAEC were cooled down on ice for 30 minutes before measuring I¹²⁵-LDL (**G**) or I¹²⁵-HDL (**H**) binding at 4°C for 60 minutes followed by radioactivity measure as above. Points in graphs represent individual experiments, bars represent the mean and error bars ± SD: *p<0.05, **p<0.01 and ***p<0.001.

The specific contribution of EL was further investigated by knocking down its expression using RNA interference. Specific siRNA against *LIPG* significantly reduced its transcript level, while the transcript levels of other genes known to limit LDL (*ACVRL1*, *LDLR*, and *SCARB1*) or HDL (*ABCG1* and *SCARB1*) trafficking remained unchanged (***Sup. Figure 1A***). The phospholipase activity in the media was also modestly but significantly reduced upon knocking down EL (***Sup. Figure 1B-C***). In the absence of EL, association of both LDL and HDL were significantly reduced (***Figure 1E-F***). After cooling the cells to 4°C to block cellular processes, binding of both LDL and HDL to HAEC was also significantly reduced in the absence of EL (***Figure 1G-H***). Together, these results support that EL facilitates the binding and uptake of both LDL and HDL by HAEC.

### 3.2 Overexpression of EL increases lipoprotein binding and uptake by endothelial cells

To corroborate the findings of the loss of function experiments and to assess the role of EL’s catalytic activity, we next overexpressed wild-type EL (WT_EL) or a catalytically inactive mutant (MUT_EL) in HAEC. Forty-eight hours post-adenovirus infection, both WT_EL and MUT_EL transcripts were significantly upregulated while the transcript levels of *SCARB1, ABCG1, ACVRL1*, and *LDLR* remained unchanged (***Sup. Figure 2A***). Correspondingly, EL protein levels were elevated following infection with both WT_EL and MUT_EL (***Sup. Figure 2B***). Upon immunofluorescence microscopy WT_EL and MUT_EL exhibited similar cellular distribution patterns with most of the signal being intracellular (***Sup. Figure 2C***). Phospholipase activity in the culture media was significantly increased upon overexpression of WT_EL but slightly decreased with MUT_EL overexpression (***Sup. Figure 2D***). Binding and association of both LDL and HDL by HAEC were significantly enhanced by overexpression of both WT_EL and MUT_EL (***Figure 2A through D***). Interestingly, we found that WT_EL showed a greater LDL association than MUT_EL. Since ¹²⁵I-lipoprotein association reflects both binding and uptake, we explored whether flow cytometry could be used to measure uptake exclusively, as cells must be detached for analysis. Endothelial cells were incubated with fluorescently labeled LDL (atto-LDL) or HDL (atto-HDL) at 4°C or 37°C. After 4 hours, the cells were detached using Accutase, and fluorescence was measured. At 4°C, the fluorescent signal was minimal and was not reduced by excess of unlabeled lipoproteins (***Sup. Figure 3A-B***). At 37°C, the fluorescent signal was detectable, and excess of unlabeled lipoproteins reduced it (***Sup. Figure 3C-D***). These findings suggest that flow cytometry is a suitable method to specifically measure lipoprotein uptake with minimal interference from membrane-bound lipoproteins. Upon overexpression of WT_EL and MUT_EL both LDL and HDL uptake were significantly upregulated with no difference whether EL was active or not (***Figure 2E-F***). All together, these results further support that EL facilitates the binding and uptake of both LDL and HDL by HAEC independently of its catalytic activity.

**Figure 2.**
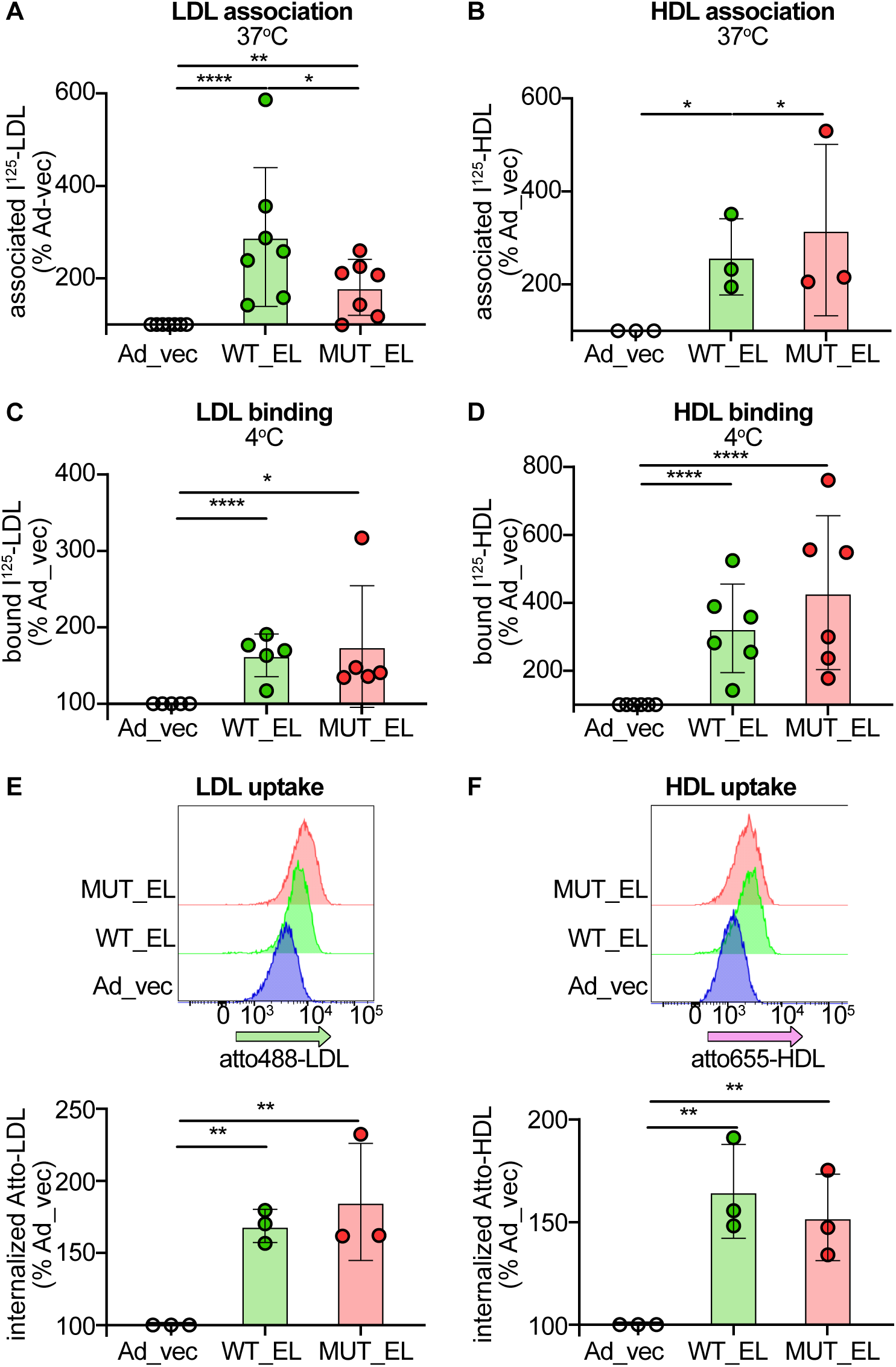
Overexpression of endothelial lipase increases LDL and HDL binding, association and uptake by human aortic endothelial cells. **A** through **D)** Two days after infection with adenovirus coding for catalytically active (EL_WT) or inactive (EL_MUT) *LIPG*, I¹²⁵-LDL (**A**) or I¹²⁵-HDL (**B**) associations or I¹²⁵-LDL (**C**) or I¹²⁵-HDL (**D**) bindings were measured as described in figure 1. (**E** and **F**) Two days after infection with adenovirus infection as described above, confluent HAEC were incubated with 50 μg/ml of Atto488-LDL or Atto655-LDL. After 3 hours, cells were washed, detached using accutase and the median intensity fluorescence was quantified by flow cytometry. Points in graphs represent individual experiments, bars represent the mean and error bars ± SD. Flow cytometry charts represent a representative experiment. *p<0.05, **p<0.01 and ***p<0.001.

### 3.3 EL catalytic activity is essential for lipoprotein transport through endothelial cells

To investigate the role of EL on lipoprotein transport through endothelial cells, HAEC were cultured on transwell systems. Because RNA interference against *LIPG* altered endothelial barrier tightness, we used pharmacological inhibitors to investigate the role of EL in lipoprotein transport. Inhibition of EL by THL significantly reduced LDL and HDL transport through HAEC (***Figure 3A-B***). Interestingly, we found that endothelial cells overexpressing WT_EL transported significantly more LDL and HDL compared to HAEC infected with an adenovirus containing an empty vector or overexpressing MUT_EL (***Figure 3C-D***). We next investigated if EL-mediated transport alters the size and lipid composition of the lipoproteins. Size exclusion chromatography of lipoproteins collected from the basolateral compartment of the transwells after transport through control and EL-overexpressing HAECs did not reveal any change in the size distributions of HDL or LDL compared to the apically applied starting material (***Figure 3E-F***). Untargeted lipidomics revealed that EL primarily increased levels of lysophophoinositol species (LPI) in LDL and HDL (***Figure 3G-H***). The same analysis did not unravel any changes of other lipids. Together these results point to minimal remodeling of the surface phospholipids of the particles without changing of the core lipids.

**Figure 3.**
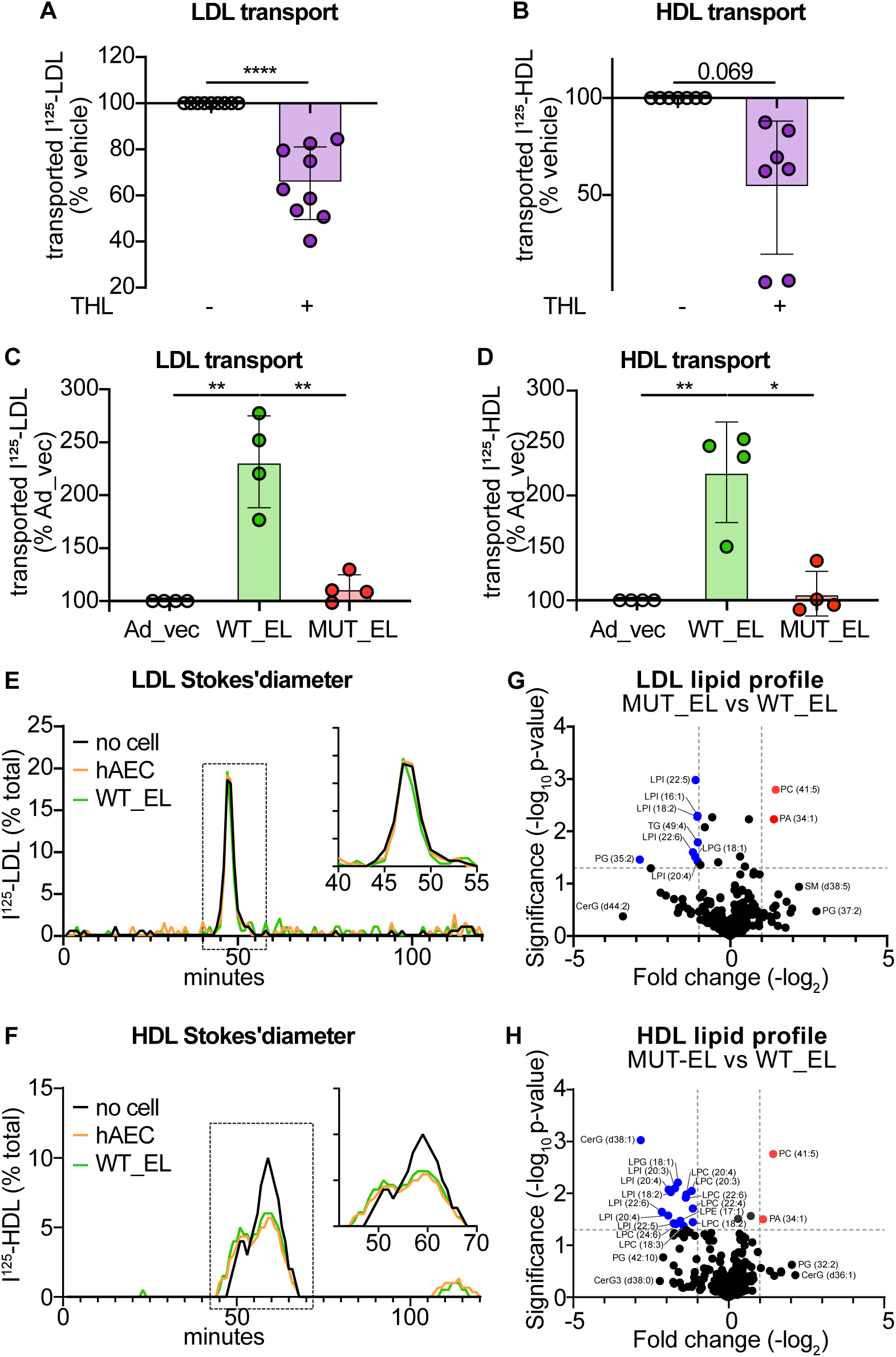
Inhibition or overexpression of endothelial lipase respectively decreases or increases LDL and HDL transport through human aortic endothelial cells. **A** through **D**) Confluent HAECs were treated with THL (A and B) or infected with adenovirus (C and D) as described in Figures 1 and 2, respectively. I^125^– LDL (A and C) or I^125^–HDL (B and D) (10 μg/mL) were added to the upper chamber of the transwell and incubated at 37 °C. After 1 hour, basolateral media were collected and radioactivity was measured using a γ-counter. **E** and **F**) Media from the basolateral chamber were loaded onto a size-exclusion column, 1 mL fractions were collected, and the radioactivity of each fraction was measured using a γ-counter. **G** and **H**) Lipid analyses of human LDL (G) and HDL (H) were performed using untargeted LC-MS/MS lipidomics. Points in graphs represent individual experiments, bars represent the mean, and error bars indicate ± SD. FPLC charts represent a representative experiment from at least two independent experiments. *p < 0.05, **p < 0.01, and ***p < 0.001.

Similarly, neither EL inhibition nor WT_EL overexpression affected the degradation of LDL or HDL (***Sup. Figure 4A through D***). In summary, these data further demonstrate that EL facilitates lipoprotein transport through HAEC without affecting their degradation.

### 3.4. ANGPTL3 reduces LDL but not HDL transport through aortic endothelial cells

We next sought to determine whether ANGPTL3 inhibits EL-dependent transport. First, using publicly available mRNA sequencing (RNAseq) data ^29^, we confirmed that HAEC do not express ANGPTL3. Next, we attempted to generate a stable cell line overexpressing ANGPTL3 using either the pcDNA3.1/Zeo-ANGPTL3 wt or the pCDNA3.1/Neo-ANGPTL3 wt plasmids; however, EAhy.926 cells transfected with ANGPTL3 were not viable. To overcome this limitation, we opted to use human recombinant ANGPTL3 produced in CHO cells and we examined whether ANGPTL3 alters LDL or HDL trafficking through HAEC. Treatment with 1 μg/ml of ANGPTL3 for 30 minutes prior adding lipoprotein significantly reduced the association and transendothelial transport of LDL but not HDL by HAEC (***Figure 4A trough D***). Afterwards, we investigated the trafficking pathways following lipoprotein uptake and aimed to determine whether EL functions as an independent receptor for LDL and HDL or if it acts sequentially with other known lipoprotein receptors on endothelial cells. We investigated the role of SR-BI, as it, like EL, has been shown to regulate both LDL and HDL binding and transport through endothelial cells ^26^. In endothelial cells lacking SR-BI, overexpression of WT_EL significantly increased HDL uptake but failed to increase LDL uptake (***Figure 4 E-F***). Interestingly the overexpression of catalytically inactive EL led to a modest but yet significantly increased LDL uptake and decreased HDL uptake compared to active EL. Together these results suggest that LDL and HDL follow different trafficking sequences with EL and SR-BI acting separately for HDL but sequentially for LDL at least in the case of active EL.

**Figure 4.**
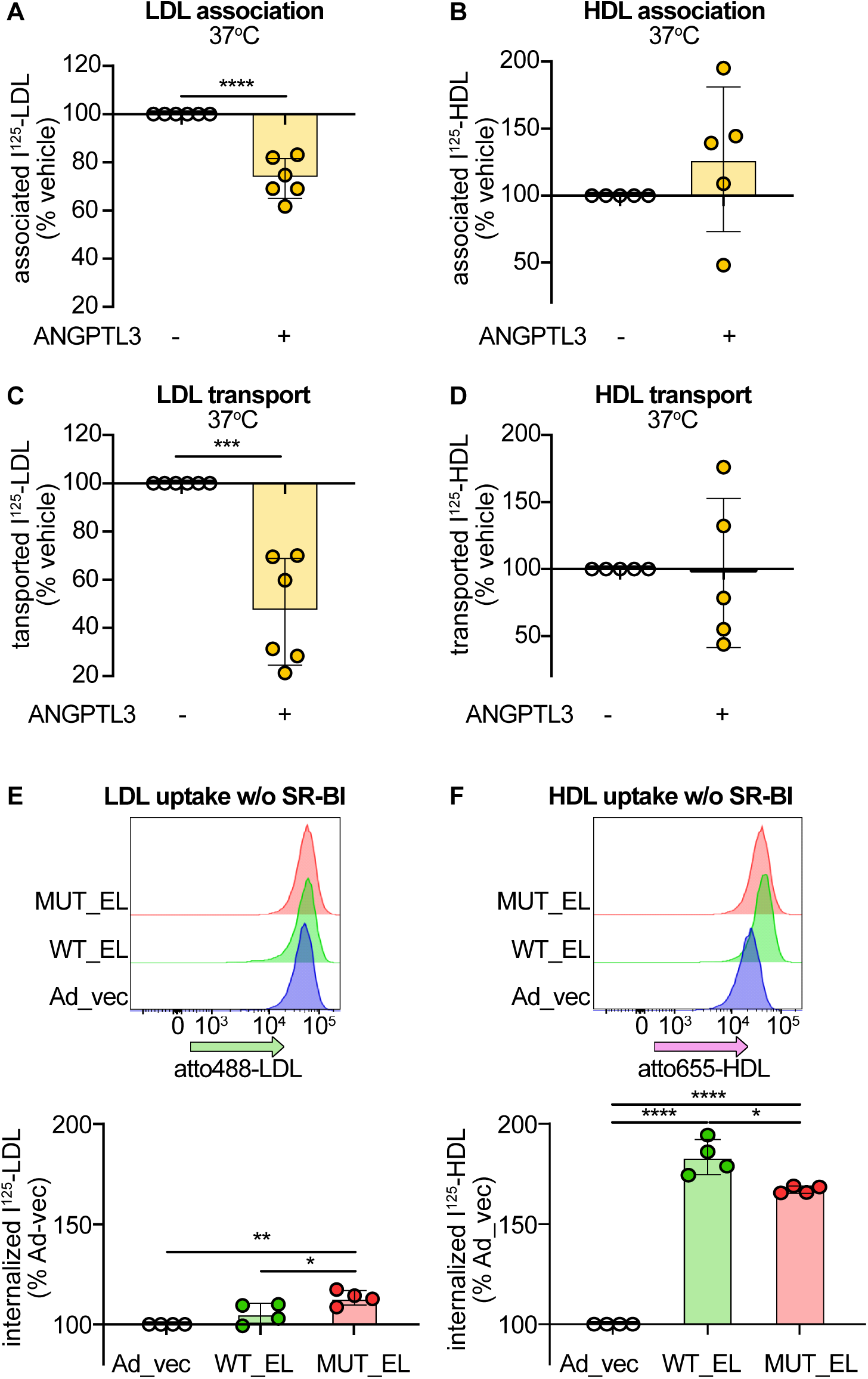
ANGPTL3 reduced LDL but not HDL transport through human aortic endothelial cells. Confluent HAEC were incubated with 1 μg/ml of ANGPTL3 for 30 minutes prior performing I^125^–LDL (**A**) and I^125^-HDL (**B**) association and I^125^-LDL (**C**) and I^125^-HDL (**D**) transport as described above. Atto488-LDL (**E**) and Atto655-HDL (**F**) association in EAhy.926 lacking SR-BI infected with adenovirus coding for EL_WT or EL_MUT were performed as described above. Points in graphs represent individual experiments, bars represent the mean, and error bars indicate ± SD. Flow cytometry charts represent a representative experiment from at least two independent experiments. *p < 0.05, **p < 0.01, ***p < 0.001 and ****p<0.0001.

We next explored whether other ANGPTL family members also limit LDL and HDL transport through HAEC. We first assessed the expression of ANGPTL4 and ANGPTL8 in HAEC using publicly available RNAseq data ^29^. While ANGPTL4 was expressed, ANGPTL8 was not expressed in HAEC ^29^. *ANGPTL4* knockdown not only failed to increase LDL or HDL association with HAEC, but even slightly reduced it (***Figure 5A-B***). Similar trend was observed in the endothelial cell line, EAhy.926 (***Figure 5C-D***). The addition of recombinant human ANGPTL8 did not change LDL and HDL association with HAECs (***Figure 5E-F***). In contrast, overexpression of ANGPTL8 in EAhy.926 cells significantly decreased the association of either lipoprotein (***Figure 5G-H***). Together, these findings highlight that ANGPTL members exhibit distinct roles not only depending on their type, but also on their endothelial origin.

**Figure 5.**
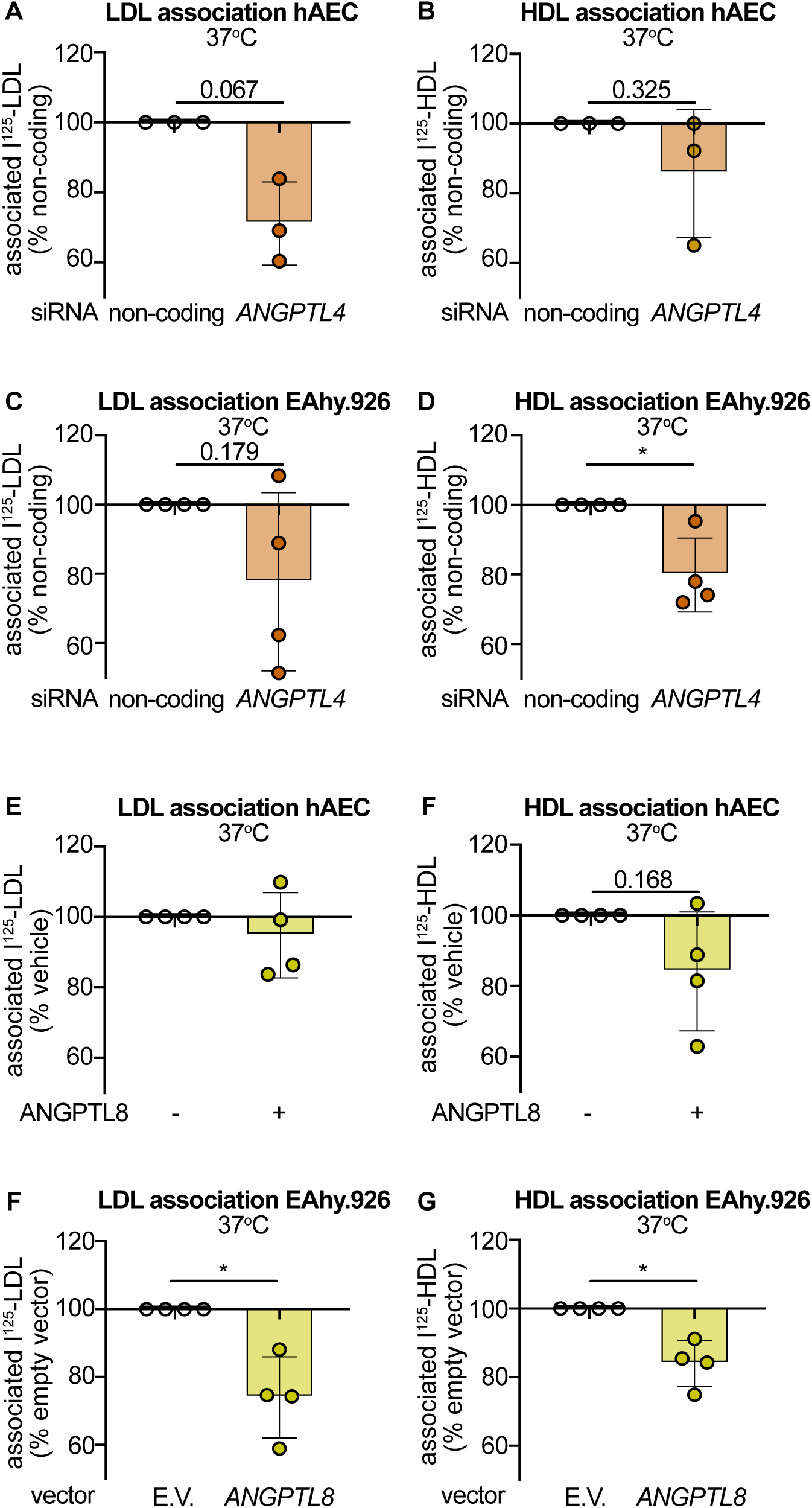
ANGPTL4 and ANGPTL8 do not reduce LDL or HDL uptake by human aortic endothelial cells. Confluent HAEC were transfected with siRNA against *ANGPTL4*. After 72 hours I^125^-LDL (**A**) and I^125^-HDL (**B**) associations were performed as described above. Confluent EAhy.926 were transfected with siRNA against *ANGPTL4*. After 72 hours I^125^-LDL (**C**) and I^125^-HDL (**D**) associations were performed as described above. Confluent HAEC were incubated with 1 μg/ml of human recombinant ANGPTL8 30 minutes prior to performing I^125^-LDL (**E**) and I^125^-HDL (**F**) associations as described above. ANGPTL8 were stably transfected into EAhy.926 cells. I^125^-LDL (**G**) and I^125^-HDL (**H**) associations were performed in confluent EAhy.926 overexpressing ANGPLT8 or empty vector control as described above. Points in graphs represent individual experiments, bars represent the mean, and error bars indicate ± SD. Flow cytometry charts represent a representative experiment from at least two independent experiments. *p < 0.05.

### 3.5. ANGPTL3 reduces LDL accumulation in bovine aorta ex-vivo

After demonstrating in a single layer of endothelial cells that ANGPTL3 regulates LDL but not HDL transport through HAEC, we explored whether EL and ANGPTL3 limit lipoprotein uptake into the bovine aortic wall. Immunostaining showed that EL is expressed not only by aortic endothelial cells but also by aortic smooth muscle cells (***Figure 6A***) in agreement with publicly available RNAseq data ^29^. To assess lipoprotein accumulation within the aorta, aortic pieces were embedded in agarose gel with only the endothelial cell layer exposed (***Figure 6B***). Following incubation with fluorescent lipoproteins, both LDL and HDL were detected within the endothelial cells (green arrows) as well as within the subendothelial layer (***Figure 6C-D***, yellow arrows).

**Figure 6.**
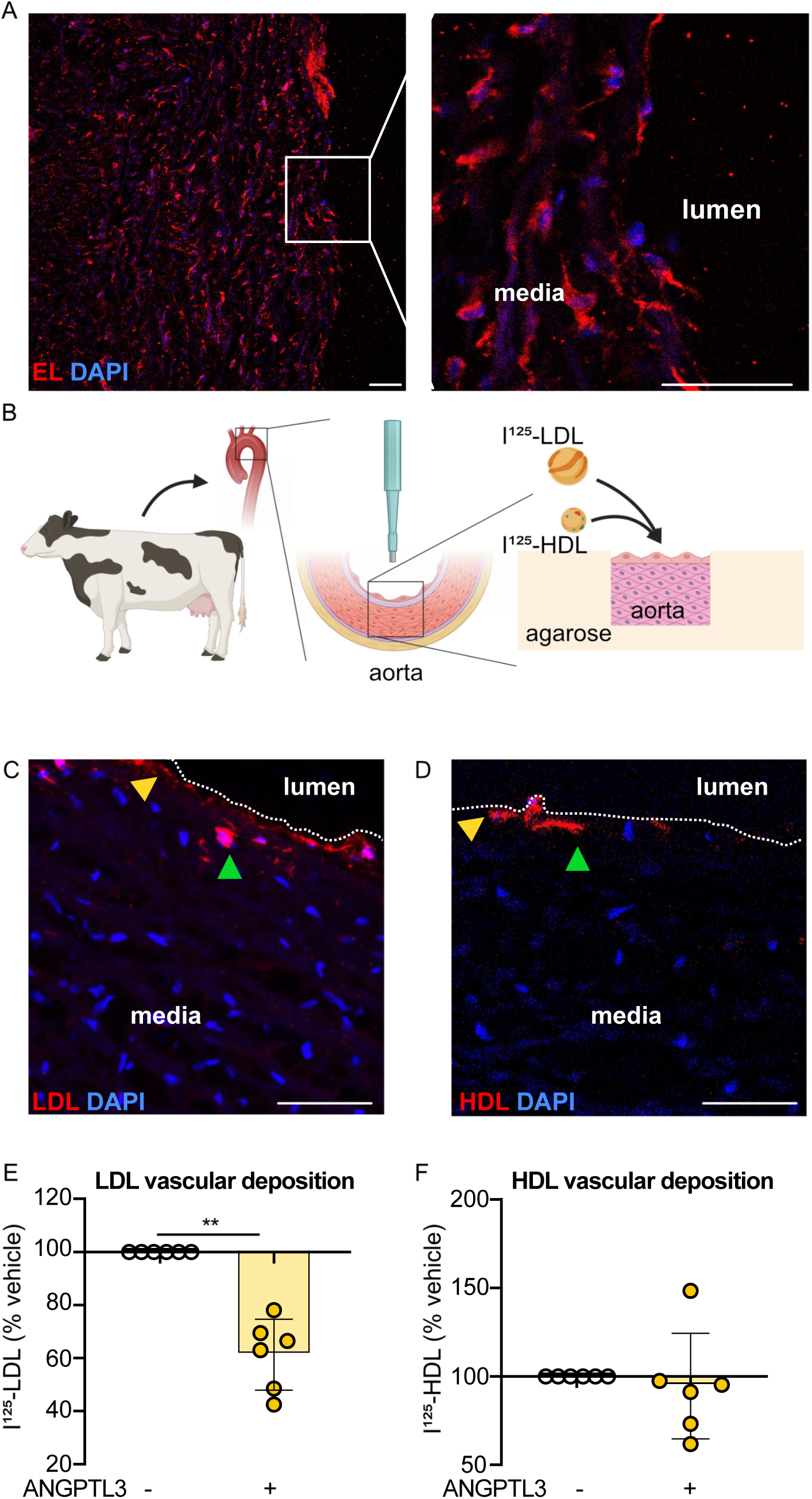
ANGPTL3 reduces LDL but not HDL deposition in aortic wall. **A**) The expression of EL (red) was assessed by immunostaining in bovine aorta. Bar: 100 μm. **B**) Schematic representation of the lipoprotein deposition within the aortic wall. **C**) Atto655-LDL (red) deposition within the arterial wall. Dashed line represents the endothelium, green arrow: endothelial deposition, yellow arrow: subendothelial deposition. **D**) Atto655-HDL (red) deposition within the arterial wall. Dashed line represents the endothelium, green arrow: endothelial deposition, yellow arrow: subendothelial deposition. **E** and **F**) Aorta were collected from the slaughterhouse, aortic punches were made and equilibrated in culture media overnight. After embedding in agarose gel, aortic punches were incubated with 1 μg/ml recombinant human ANGPTL3 for 30 minutes followed by incubation with 10 μg/ml of I^125^-LDL (E) or I^125^-HDL (F). After another hour at 37°C, punches were extensively washed and radioactivity was counted using a γ-counter. Points in graphs represent individual aorta, bars represent the mean, and error bars indicate ± SD. Microscopy images are representative of at least two individual aorta. Bars: 100 μm. **p < 0.01

Next, we evaluated the effect of ANGPTL3 by pre-treating the aortic pieces for 30 minutes before adding the lipoproteins. After one-hour incubation with I^125^–LDL or I^125^– HDL, ANGPTL3 treatment significantly reduced LDL accumulation (***Figure 6E***), while HDL accumulation was not affected (***Figure 6F***).

## 4 Discussion

The role of ANGPTL3 has been preliminarily investigated with a focus on plasma lipoprotein concentrations. In the liver, ANGPTL3 inhibits the LDLR independent uptake and removal of LDL by EL. In addition and together with ANGPTL8, ANGPTL3 inhibits lipoprotein lipase, which hydrolyses triglycerides in VLDL ^30^. As the result, ANGPTL3 inhibition by antibodies or siRNAs has been developed for the treatment of homozygous familial hypercholesterolemia and mixed hyperlipidemia, respectively ^5–7,10,13,14^. While elevated levels of LDL and VLDL remnants are accepted as causal risk factors of ASCVD, it is the accumulation of LDL in the arterial wall after transendothelial transport that directly promotes atherogenesis ^20^. HDL-mediated efflux of cholesterol from macrophages of the arterial wall for reverse transport to the liver requires transendothelial transport of HDL for both the entry and the exit ^20^. Since EL is mainly expressed in endothelial cells, we investigated if EL and its inhibitor ANGPTL3 contribute to the trans-endothelial transport of LDL and HDL.

We found that EL, which is bound to endothelial cell surface promotes LDL binding, uptake, and trans-endothelial transport in aortic endothelial cells, with ANGPTL3 further modulating these effects. These results suggest that the beneficial effects of ANGPTL3 inhibition on plasma concentrations of LDL may be counterbalanced by increased LDL deposition in the arterial wall and explain the unchanged ASCVD risk of individuals carrying LOF variants of ANGPTL3 despite their lower levels of LDL-C ^18,19,31–33^. Importantly, while preclinical models lacking ANGPTL3 have shown a reduction in atherosclerotic lesions ^5^, this finding contrasts with our results and may be attributable to ANGPTL3’s inhibitory effect on LPL ^34^. In particular as *LPL* in contrary to *LIPG* LOF mutations are associated with a significantly increased risk of coronary artery disease (CAD) ^19,33^. Further, a recent study showed that the level of ANGPTL3 alone showed only a modest positive association with lipoprotein concentration and no association with coronary heart disease (CHD) while the concentration of the complex ANGPTL3/8 in the plasma, which strongly inhibits LPL, is associated with atherogenic lipid profile and CHD ^18^. The role of LPL in lipoprotein transport was not investigated here, as aortic endothelial cells do not express it ^29^. However, LPL has previously been shown to enhance LDL uptake by mouse aortic endothelial cells via LDLR, which induced LDL degradation in the absence of inflammation ^35^. Additionally, reduced atherosclerotic plaque size in *Angptl3^-/-^* mice has been linked to decreased monocyte infiltration and inflammation ^36^. Notably, lipid deposition in *Angptl3^-/-^* models was not specifically examined ^5,36^, leaving open the possibility that lipid accumulation might have increased, but reduced monocyte infiltration and inflammation led to plaque reduction. This hypothesis warrants further investigation.

In addition to targeting ANGPTL3, blocking EL has been proposed as another therapeutic approach, currently under investigation in clinical trials ^37,38^. However, preclinical findings on EL are inconclusive: *Lipg^-/-^;apoE^-/-^* double knockout mice exhibit significantly fewer atherosclerotic lesions ^39^, while other studies report no effect on atherosclerosis development on *Apoe^-/-^* or *Ldlr^-/-^* backgrounds when EL is absent ^40^. Nonetheless, patients with EL LOF mutations show a reduced risk of CAD despite elevated LDL-C plasma levels ^16,31^. Conversely, EL gain-of-function may increase CAD risk despite lower LDL and triglyceride levels ^31^. These opposing outcomes are thought to be mediated by changes in HDL-C levels, as HDL without EL contact has been shown to have increased cholesterol efflux capacity ^16,37^. Contradicting these findings, HDL exposed to EL has demonstrated enhanced antioxidative capacity, anti-inflammatory activity, and promoted nitric oxide (NO) production ^25,41,42^. Furthermore, we here confirmed our previous finding that EL limits HDL transport through aortic endothelial cells, a critical step preceding cholesterol efflux from subendothelial macrophages ^22^. Our findings present a novel perspective on anti-atherogenic strategies: blocking EL might also reduce atherosclerosis by limiting LDL entry into the arterial wall.

Interestingly, our current study shows that while EL regulates the transport of both LDL and HDL through endothelial cells, ANGPTL3 specifically reduces LDL transport. This difference may be due to several factors. First, we demonstrated that EL-mediated LDL transport but not EL-mediated HDL-transport depends on SR-BI. This is consistent with our earlier observations that activation of sphingosine-1-phosphate receptors (S1PRs) increases HDL transport via SR-BI regulation on the cell membrane while reducing LDL transport ^43^. Our finding that SR-BI and EL interact in LDL but not HDL transport may explain this discrepancy. Second, studies have detected ANGPTL3 as a component of HDL in humans and mice ^44,45^. Interestingly, these studies found reduced ANGPTL3 levels in diabetic patients, who have a higher prevalence of atherosclerosis. Although we did not measure ANGPTL3 levels in HDL in our study, its presence might be sufficient to inhibit the EL pathway. Third, while ANGPTL3 is present in HDL, Kraaijenhof and colleagues found that ANGPTL3 is more active when bound to LDL ^46^.

In addition to ANGPTL3, we investigated the roles of two other ANGPTL family members: ANGPTL4 and ANGPTL8, which are also in consideration as targets of lipid lowering therapeutics ^47–49^. Our findings confirmed that ANGPTL4 does not inhibit EL, because knocking down ANGPTL4 expression did not increase but rather decreased LDL or HDL uptake by endothelial cells. The reduction of LDL transport through HAEC in the absence of ANGPTL4 might also partially explain the protective effect observed in *ANGPTL4* LOF mutations ^19,31,32^. In contrast, ANGPTL8 had no effect in HAEC but reduced both LDL and HDL uptake in EAhy.926, suggesting a different mechanism of action in cells of different endothelial origin, which remains to be investigated. Recent findings showed that ANGPLT8 is required to form a dimer with ANGPTL3 to inactivate EL ^30,50,51^. This observation may explain the difference in effect magnitude between ANGPTL3 and ANGPLT8 however we did not investigated here the role of ANGPTL3 and ANGPTL8 dimers.

Our research has several limitations. Primarily, we relied on cell culture models to investigate the roles of EL and ANGPTL3, motivated by the desire to isolate the effects on the aorta from those on the liver. However, to address this limitation, we developed a novel *ex vivo* model using bovine aorta to study lipoprotein accumulation in the vascular wall. In this study, we used recombinant human ANGPTL3 because endothelial cells transfected with ANGPTL3 vectors did not proliferate. These findings are consistent with previous reports that ANGPTL3 can promote apoptosis ^52^. However, previous studies also used recombinant human ANGPTL3 at similar concentration to investigate different questions ^36,53^.

In conclusion, our study demonstrates that both EL and its plasma inhibitor ANGPTL3 regulate the deposition of LDL within the vascular wall. These findings may have significant implications for the development of ANGPTL3 inhibitors, and we recommend that atherosclerotic plaque volume should be closely monitored in clinical trials. Notably, in the context of homozygous familial hypercholesterolemia (hoFH), the hepatic benefits of ANGPTL3 inhibition may outweigh the risks of lipoprotein deposition within the arterial wall, especially as five-year follow-up studies have shown no increased mortality in hoFH patients taking evanicumab ^54^. However, this benefit may not occur in other patients whose LDL receptor activity can be normalized or even exaggerated by treatment with statins, bempedoic acid, ezetimibe, or PCSK9 inhibitors.

## 5. Acknowledgements

JR and AvE were supported by a grant from the Swiss Heart Foundation (No. FF22028). JR was further supported by the Synapsis Foundation (No. 2022-PI04), a Swiss lipid Award from the AGLA and an Alzeimer’s disease Award from the BrightFocus foundation (No. A2021037S). AvE was also supported by a grant from the Swiss National Science Foundation (No. 31003A-160216). MAL is supported by grant from EMPIRIS Foundation, Zürich.

The authors wish to thank Paul Müller, Lukas Fuchs, Yannick Stichamd, and Harald Manfred at the slaughterhouse of the Vetsuisse Faculty University of Zurich for their help providing the bovine aortas.

## 7 Participation

JR and AvE conceived the project, GP, WEIT, EV, JR, MAL, ES, and SB, generated and analyzed the data, SF generated and provided the viruses, LR and KF generated and characterized the shRNA cells. TL provided bovine tissues. JR and AvE acquired the funding. JR generated the first draft of the manuscript. All authors reviewed, edited and agreed on the final manuscript

## 8 Figure legends

**Supplemental Figure 1. SiRNA against LIPG reduced specifically transcript expression and phospholipase activity. A**) Expression of *LIPG*, *SCARB1*, *ABCG1*, *ACVRL1*, *LDLR, AP2M* and *CAV1* transcripts were measured by real time quantitative RT-PCR and normalized to the expression of Actin three days after transfection. **B**) Phospholipase activity was measured in media 72 hours after siRNA transfection with phospholipase A1 substrate. Links graph is a representative experiment and right graph is the quantification of the area under the curve of four different experiments. Points in graphs represent individual experiment, bars represent the mean, and error bars indicate ± SD. *p < 0.05, ***p<0.001.

**Supplemental Figure 2. Infection with adenovirus coding for wild type EL increased specifically transcript expression and phospholipase activity. A**) Expression of *LIPG*, *SCARB1*, *ABCG1*, *ACVRL1*, *LDLR, AP2M* and *CAV1* transcripts were measured by real time quantitative RT-PCR and normalized to the expression of Actin 48 hours after the infection with adenovirus coding for catalytically active (EL_WT) or inactive (EL_MUT) *LIPG*. **B**) EL protein level was measured by Western blot 48 hours the infection. **C**) EL (red) cellular localisation was assessed by immune staining 48 hours the infection. bar: 50 μm **D**) Phospholipase activity was measured in media 72 hours after siRNA transfection with phospholipase A1 substrate. Links graph is a representative experiment and right graph is the quantification of the area under the curve of three different experiments. Points in graphs represent individual experiment, bars represent the mean, and error bars indicate ± SD. ***p<0.001.

**Supplemental figure 3. The uptake of lipoproteins is measured by flow cytometry.** Confluent cells were incubated at either 4°C or 37°C with 50 μg/ml of Atto655-LDL or Atto655-HDL in the absence (blue) or presence (orange) of 100x respective non-labeled lipoprotein. After 3 hours, cells were detached with accutase and median fluorescence was recorded by flow cytometry.

**Supplemental figure 4. Absence or overexpression of endothelial lipase do not alter LDL and HDL degradation by human aortic endothelial cells.** After RNA silencing or infection with adenovirus as described in figure 1 and 2 respectively, confluent HAEC were incubated with 10 μg/ml of I^125^-LDL or I^125^-HDL. After 4 hours, media was collected and degradation products were isolated after acid precipitation and chloroform extraction and compared to the associated respective lipoprotein determined as above. The Percentage of degradation per association was calculated by dividing the cpm of degradation by the sum of association cpm + degradation cpm * 100. Points in graphs represent individual experiment, bars represent the mean, and error bars indicate ± SD.

