## Supplementary figures and images for "Angiopoietin like protein 3 regulates low-density lipoprotein transport through aortic endothelial cells via endothelial lipase"

### Sup figure 1

**Supplemental Figure 1**

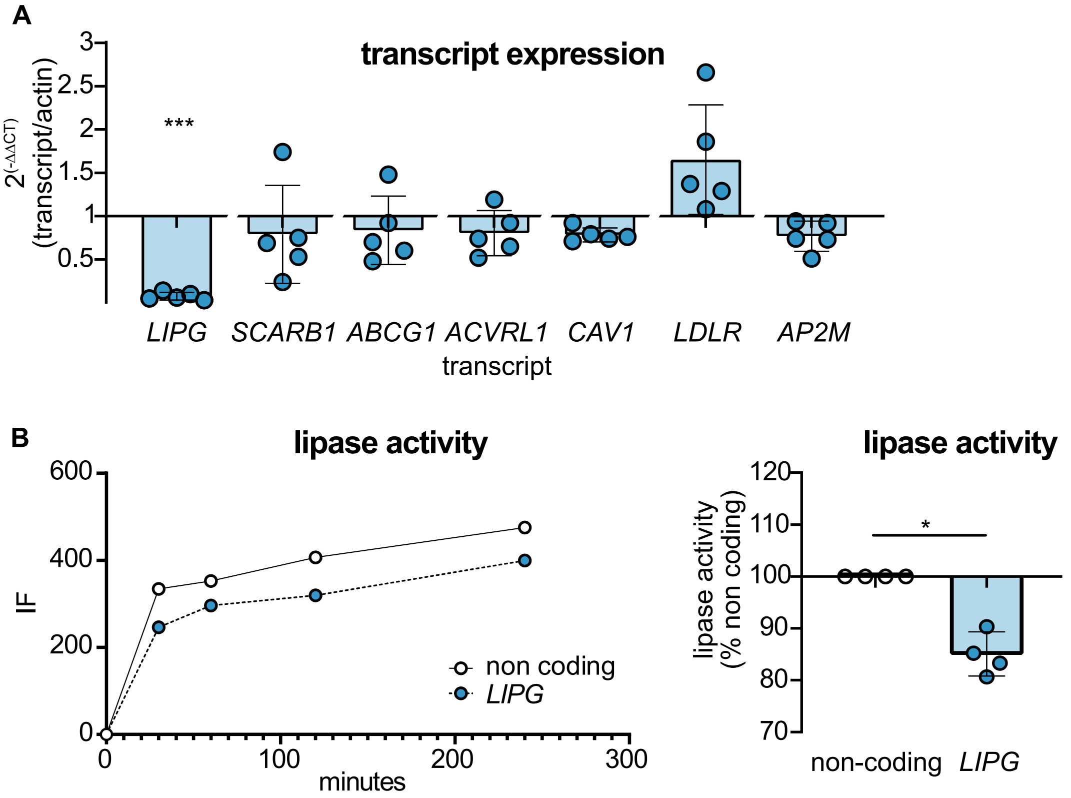

### Sup figure 2

**Supplemental Figure 2**

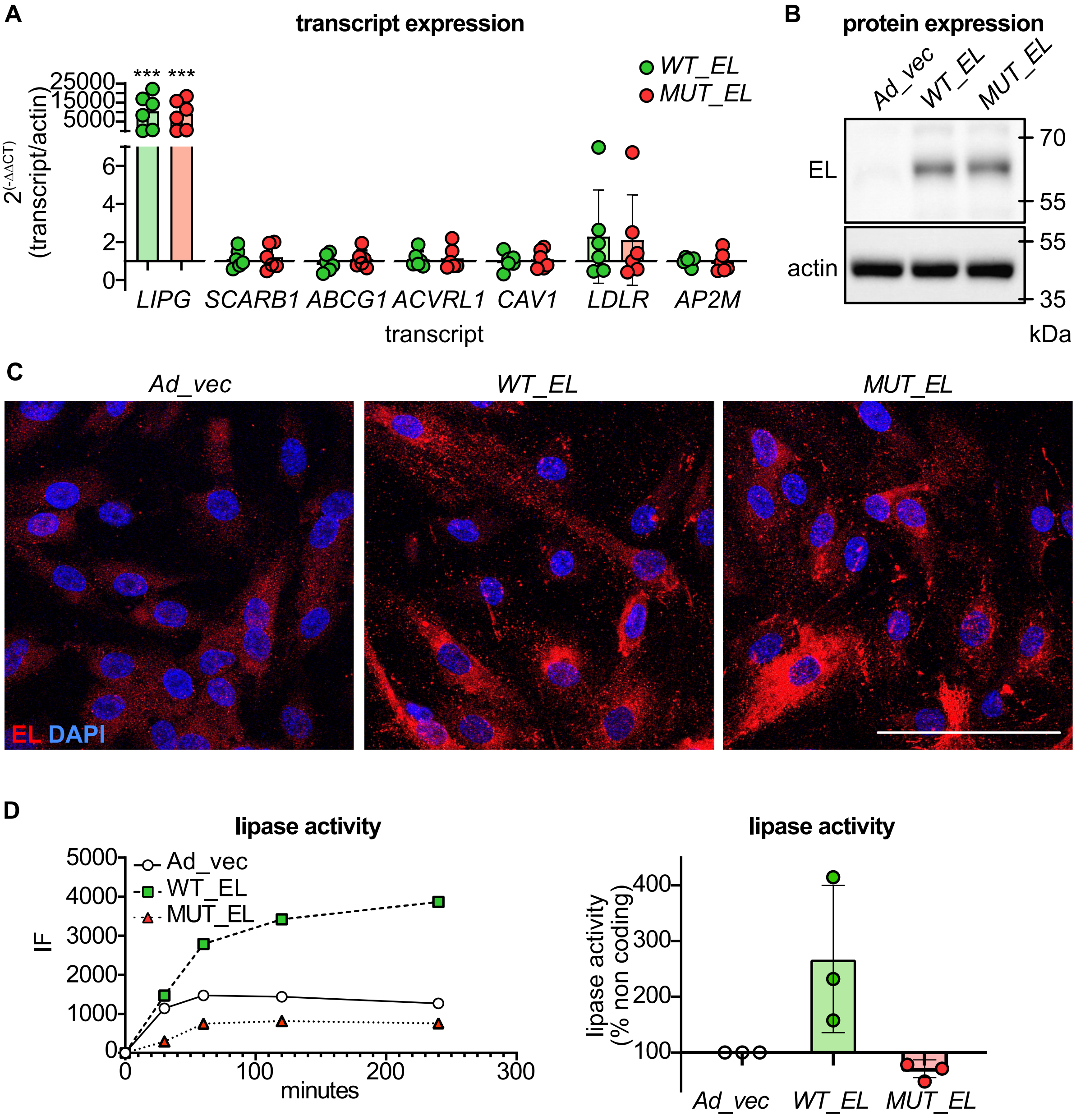

### Sup figure 3

**Supplemental Figure 3**

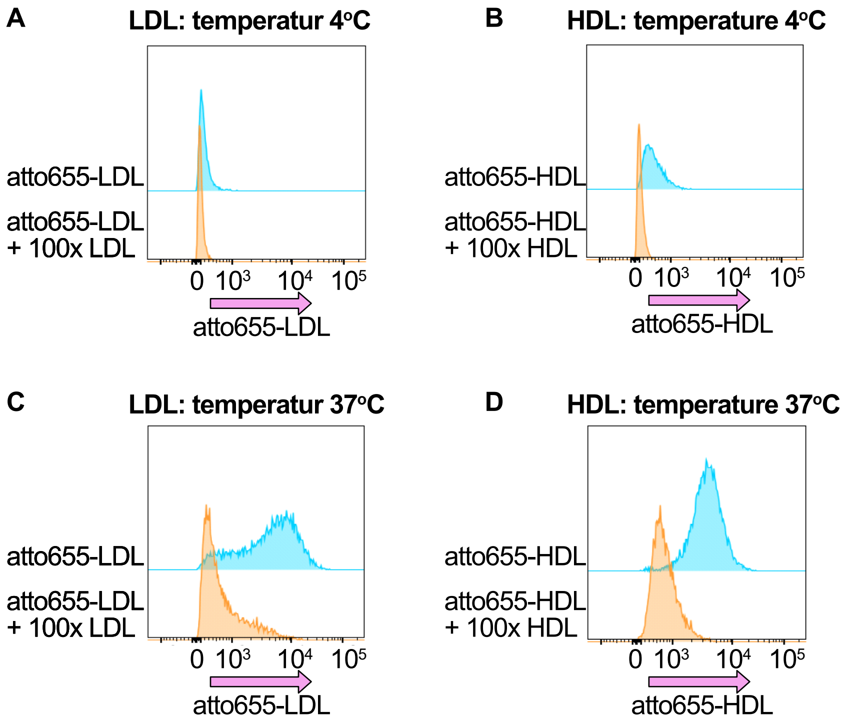

### Sup figure 4

**Supplemental Figure 4**

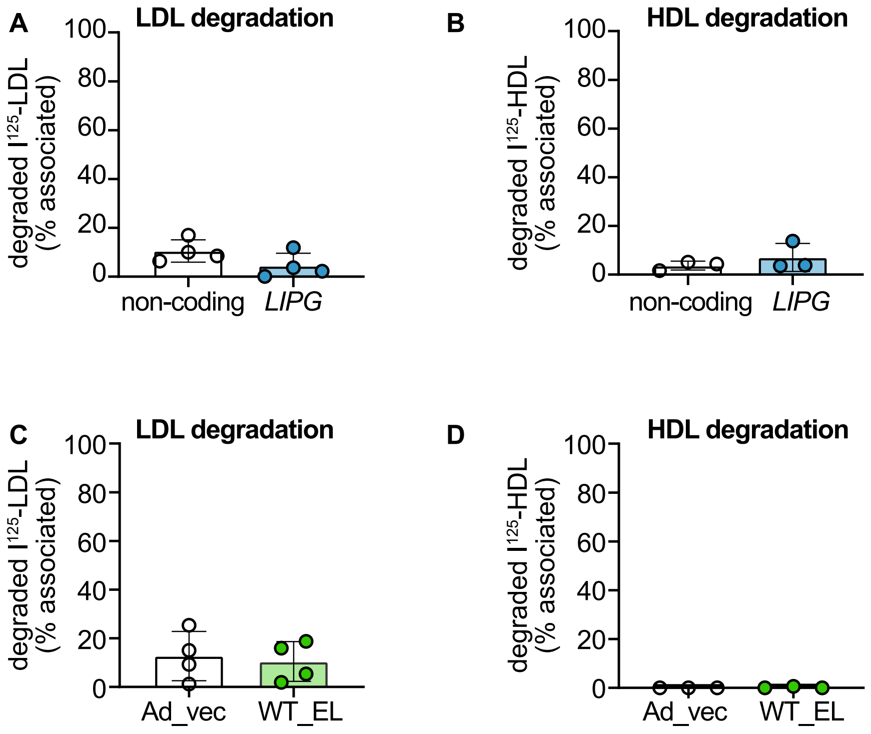
